# High-Spatiotemporal Imaging of Protein Secretion During Cell-to-Cell Communication via Integrative Biosensing Nanoplasmonic Array

**DOI:** 10.1101/2025.05.17.654581

**Authors:** Younggeun Park, Partha Ray, Katsuo Kurabayashi

## Abstract

Protein secretion underpins diverse physiological processes in cell-to-cell communication, tissue homeostasis, and the onset of diseases. Mapping the secretomes from paired cells provides avenues for understanding their interactions. However, prevailing approaches yield only semi-quantitative endpoint data, lacking real-time and quantitative information. Here, we present real time spatiotemporal imaging of extracellular secretions from individual cells via a high-throughput integrative biosensing nanoplasmonic array (iBNA) with a microfluidic chamber. The self-assembled iBNA, composed of precisely arranged gold nanostructures and functionalized with aptamer receptors, enhances plasmonic resonance and significantly improves the spatiotemporal resolution and specificity of interleukin-6 (IL-6) imaging, surpassing gold-standard techniques. The molecular recognition of iBNA, and sensing mechanism exploits biomolecular surface binding-induced localized plasmonic resonance shifts, correlating with cytokine concentration and enabling optoelectronic detection of the transmitted light. Using this approach, we achieve spatiotemporally resolved visualization of IL-6 secretion dynamics at the single-cell level and unveil the temporal and polarized variation of cell-cell communications. This transformative platform holds significant potential to advance immunological research, cellular biology, and diagnostic applications for infectious diseases by enabling unprecedented insights into the spatiotemporal patterns of protein secretion in individual cells.

## 1. Introduction

The immune system’s ability to defend against endogenous threats, such as cancer cells and exogenous pathogens, including bacteria, viruses, and fungi, requires highly sophisticated regulatory mechanisms, much of which remain incompletely understood.[1–3] A key contributor to the immune system’s complexity is the dynamic nature of interactions between immune cells over time. Immune cells, fundamental to immune defense, are diverse subsets distinguished by their functional responses to pathogens or pathogenic molecules, such as endotoxins. Understanding these immune cells’ intricate and evolving functional activities is crucial to developing predictive models of immune responses in human diseases. [1,2,4–6]

Typically, immune cell functionality is characterized by the patterns of protein secretion, with cytokines being the most notable.[7–10] Cytokines are vital, water-soluble signaling proteins that regulate immune cell populations’ development, proliferation, and functionality and mediate intercellular communication.[9,11] Notably, cytokine secretion varies within cell populations, reflecting a phenomenon known as cell heterogeneity. For instance, T cells, indispensable players in immunity, are classified in subsets such as regulatory, helper, and cytotoxic cells. Despite similarities in morphology, size, and surface markers, these subsets exhibit differential protein expression depending on environmental stimuli or specific activation contexts.[12–16]

Analyzing the cytokine secretion patterns of immune cells from blood, lymph nodes, or tissues provides valuable insights into human inflammatory immune responses.[9,15,15] Accurate, real-time monitoring of these cytokine production patterns is essential to comprehensively evaluate therapeutic interventions for allergies, immunodeficiencies, transplant rejection, asthma, autoimmunity, and infections. Elucidating the full spectrum of immune system functions at the single-cell level requires precise resolution.

Fluorescence imaging is the gold standard, enabling live-cell imaging that captures the temporal evolution of cytokine intracellular localization and transcriptional processes initiated by intercellular communication.[17–21] An innovative advancement has been the integration of optical microscopy with flow cytometry, which augments cytometry by introducing two-dimensional imaging capabilities. This arrangement introduces two-dimensional imaging capability to cytometry, which provides several new data dimensions, including cellular morphology[22,23] cell-signaling induced nuclear localization of transcription factors,[21,24,25] protein localization to immune synapses,[24,26] and cellular update of fluorescent particles.[27,28]

Despite the significant contributions of these methods to scientific knowledge, they predominantly offer indirect insights into intercellular communication, as they rely on labeling agents and fail to observe the dynamic, real-time interactions occurring between cells directly. To address this limitation, photonic resonant imaging has been employed to map signaling protein secretion from individual cells and model cell adsorption kinetics. However, this approach has drawbacks, including the inability to quantify single-cell secretion levels, limited experiment duration (up to 2 hours), and low throughput (analyzing only a few dozen cells). Furthermore, the inadequate probe resolution prevents molecular-scale visualization of cytokine distribution, which is critical for understanding cell-cell communication. Current methodologies fail to provide real-time, high-resolution imaging of cytokine release and propagation patterns during intercellular communication.

In this study, we introduce an innovative approach for direct, high-resolution mapping of cytokine-mediated intercellular communication, encompassing the release and reception of signaling proteins between individual cells, utilizing an integrated biosensing nanoplasmonic array (iBNA) constructed from a self-assembled plasmonic nanostructure array functionalized with aptamer receptors. The term "bioinspired nanoplasmonic array" refers to an innovative system that leverages refined biological recognition mechanisms to enhance nanoscale optical performance. It consists of a densely arranged and uniformly distributed array of nanoplasmonic structures functionalized with aptamers. Aptamers are nucleic acid ligands that bind to their cognate protein molecules with high sensitivity and specificity, comparable to or superior to monoclonal antibodies. Moreover, cheaper manufacturing costs and longer storage life at room temperature make them more appealing for diagnostic and analytical applications than antibodies.[29] Additionally, due to their smaller molecular size and dimensions, aptamers (6-30 kDa, 2 nm) deliver superior resolution and less steric hindrance than antibodies (150-180 kDa, 15 nm) during molecular-scale interactions.[30] Thus, this system achieves highly sensitive and selective imaging for various applications by combining aptamers’ precise molecular recognition properties with the advanced optical capabilities of nanoplasmonic arrays. We achieve significantly enhanced sensitivity and specificity for IL-6 detection—100 times greater than traditional methods. The imaging technique exploits localized surface plasmonic resonance (LSPR) changes induced by biomolecular binding events, modulating incident light delivery via iBNA. The concentration of cytokines secreted onto the iBNA dictates the LSPR tuning, which is monitored optoelectronically. Leveraging the highly sensitive and dynamically adaptive imaging platform, we enable real-time, high-resolution spatial imaging to precisely characterize IL-6 secretion dynamics at the live single-cell level. By simulating the dynamic secretion profiles of IL-6, we visualize cell-cell communication dynamics between Jurkat T cells and CD4+ T cells under varying spatial arrangements.

## 2. Results and Discussions

### 2.1. Principle of iBNA for cytokine imaging

Figure 1 shows the underlying concept of iBNA for cytokine imaging. The surface of iBNA on the SiO2 layer is conjugated with aptamers that selectively target cytokine IL-6. Without the targeted cytokine molecules, the aptamer-conjugated iBNA collects incident light at λ = ∼532 nm due to the LSPR effect (“OFF” mode). In the OFF mode, light transmission through the SiO2 layer becomes low, keeping most incident light from reaching the light-sensitive sensor. The binding of IL-6 molecules onto the aptamer-conjugated iBNA surfaces shifts the plasmonic resonance wavelength owing to a change in the local refractive index (RI) near the iBNA surfaces.[31] A larger portion of the incident photons then transmits through the thin SiO2 layer and reaches the underlying light-sensitive sensor (“ON” mode). The ON mode results in a redshift of the extinction spectrum peak of the iBNA, thus leading to an increased detection signal. The detection intensity is determined by the cytokine concentration of a sample solution deposited on the device surface covered with the iBNA layer. Obtaining a correlation between the photocurrent change and the cytokine concentration allows highly sensitive and spatial quantification of IL-6.

**Figure 1.**
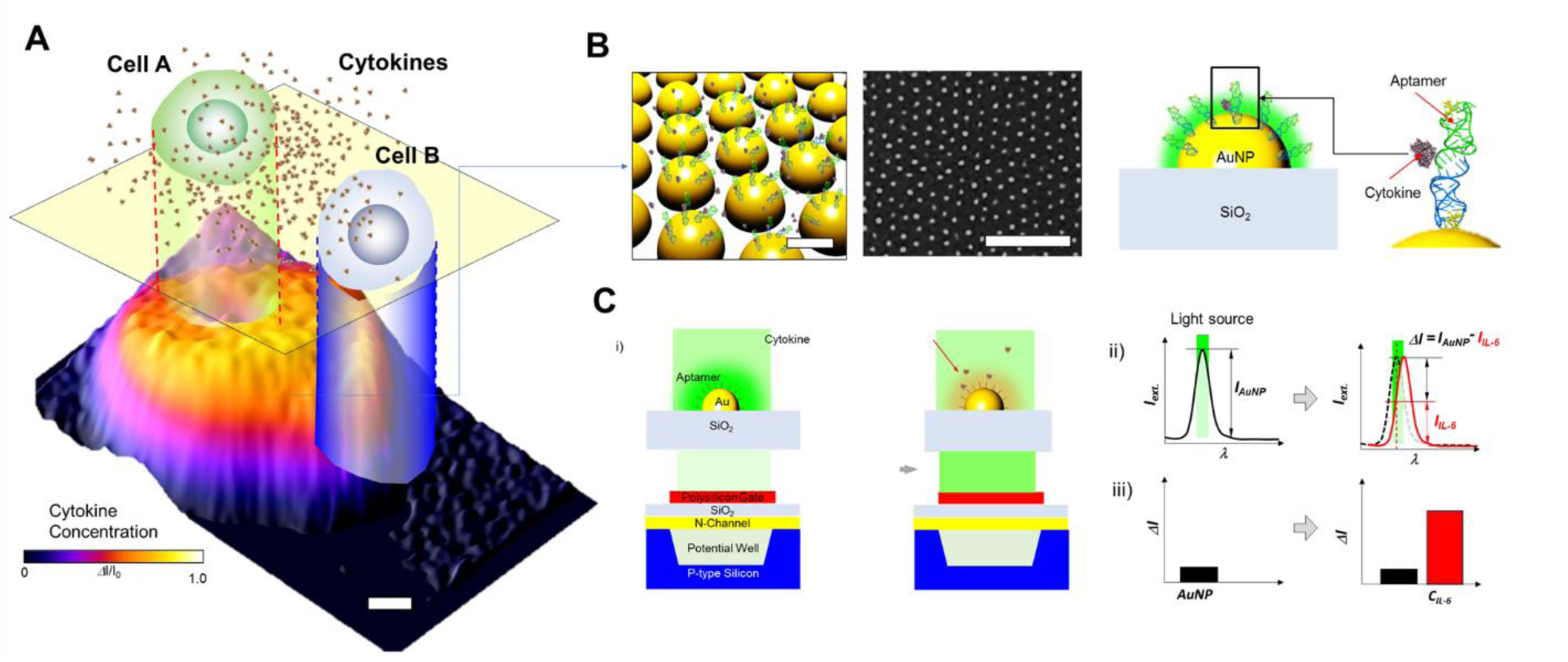
High spatiotemporal imaging of protein secretions from single cells via iBNA. (A) iBNA enables high spatiotemporal imaging of cytokine secretion amid cell-to-cell communication at the single level, (B) iBNA consists of uniform and high-density probe structure arrays functionalized with aptamer receptors, ensuring to capture a specific protein secreted by the cells to the other cells. (C) Schematic of iBNA for cytokine imaging. i) An aptamer-attached iBNA (d = 50 nm) resonates with incident light at λ = 532 nm. The resonance induces strong extinction around the Ap-iBNA on SiO2. Due to the strong extinction, the limited power density of incident light is delivered to the charge-coupled device (CCD)—the plasmonic resonance between the iBNA and the incident light results in a lower photo intensity signal. When cytokines selectively bind to the aptamers on the iBNA, there is a change in the local RI. These local refractive index changes decrease resonance between the cytokine-Ap-iBNA and the incident light. A larger portion of light-power density can be delivered to the CCD image sensor. ii) The extinction of the iBNA is matched to the incident light wavelength. Local RI change based on cytokines binding on the aptamer-iBNA leads to a shift of the extinction peak. The extinction peak and wavelength of the incident light are no longer matched. iii) Resonance between the plasmonic extinction of the aptamer-AuNP and the incident light source induces a decrease in the incident light detected in CCD. The extinction peak shift leads to more incident lights being detected.

Applying the nanoscopic imaging principle of IL-6 molecules on a large scale that can cover a higher-than-cellular level, uniform and dense plasmonic structure of iBNA in a large area is essential. The low density of sensing probes leads to the low spatial resolution of IL-6 imaging— poor uniformity of sensing probes on the substrate results in weak and broad extinction and poor spectral specificity.

Therefore, constructing a high-density and uniform plasmonic nanostructure layer is necessary to achieve a highly sensitive large-area image—a uniform distribution of AuNPs results in a narrow LSPR spectrum curve. Conventional fabrication methods, including drop casting deposit and E-beam deposition, have been employed; however, they are associated with poor uniformity or low density. [31–33] To address this issue, we structurally engineered self-assembled AuNP arrays with an optimal interparticle distance and density.

### 2.2. Uniform and dense nanoplasmonic structure array

Figure 2 shows the construction that enables fine imaging resolution and a uniform and dense array of plasmonic nanostructures. The major geometric parameters of iBNA that affect the field of local surface plasmon resonance and the harmonic effects on the resonance and sensitivity are the densities and the interparticle distance (*linter*) of plasmonic nanostructures (Figure 2A).

**Figure 2.**
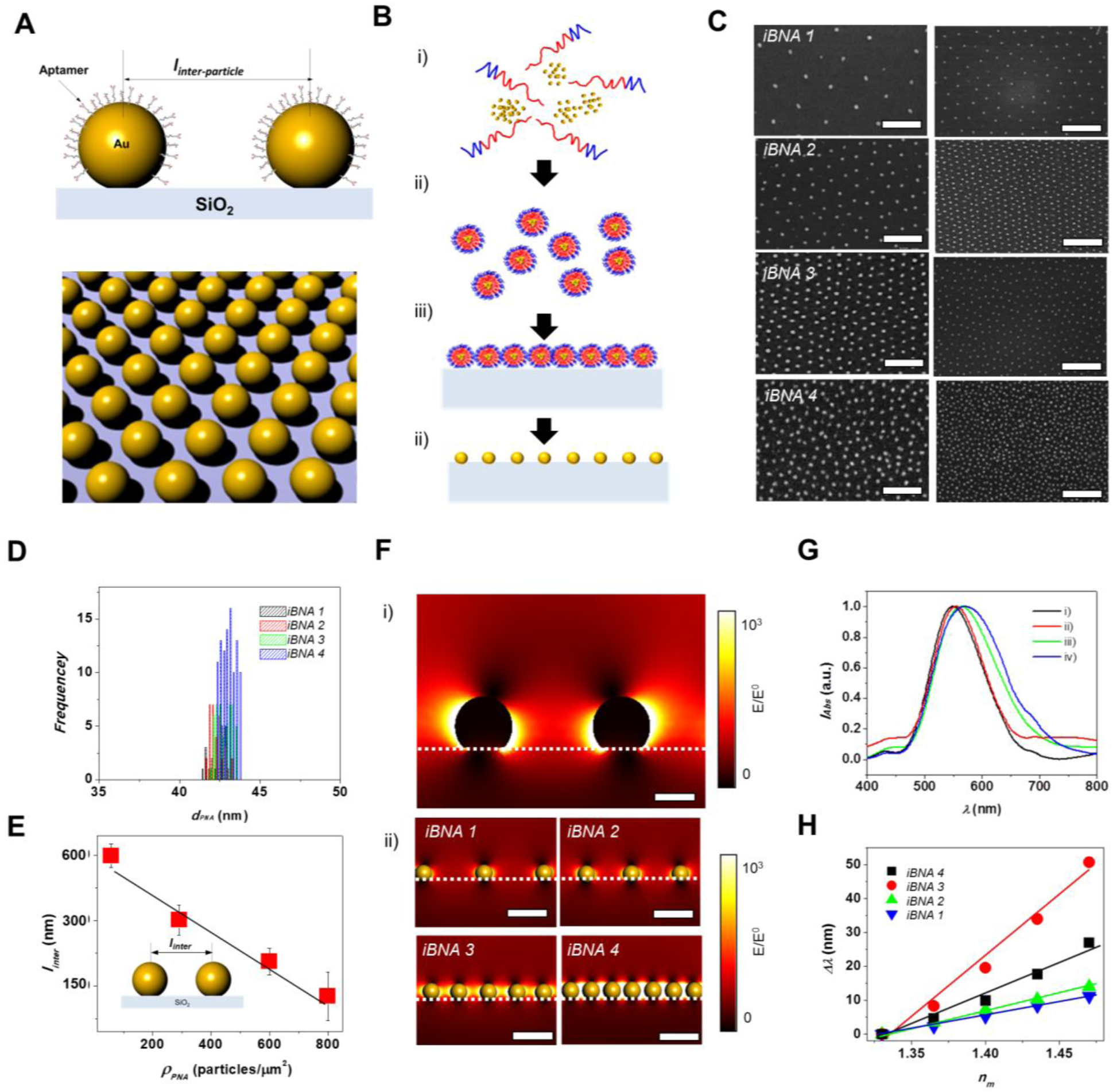
Construction and characterization of iBNA. A) Schematics of plasmonic nanostructure array and inter-nanostructure distance, linter. B) Schematics of the iBNA fabrication steps: i) self-assembly of block copolymer and gold precursor, ii) formation of nano-sized building block structure, iii) layering of the nano-sized building blocks on an optically transparent substrate, and iv) removal of organic parts of the nano-sized building blocks and reduction of gold ions to form the nano metallic structure. C) scanning electron microscopy (SEM) images of iBNA at i) 75, ii) 250, iii) 600, iv) 800 particles/μm2 (scale bars = 500 nm and 1000 nm); D) size distribution of iBNA as a function of particle density from 75 to 800 particles/μm2. E) linter, of iBNA as a function of particle density from 75 to 800 particles/μm2. F) Electric field intensity distributions around two AuNP structures at i) linter = 150 and ii) of iBNA 1 (linter = 600 nm), iBNA 2 (linter = 300 nm), iBNA 3 (linter = 100 nm), and iBNA 4 (linter = 75 nm). G) extinction spectra of i) iBNA1, ii) iBNA2, iii) iBNA3, and iv) iBNA4. H) LSPR peak shift versus refractive index (RI) for samples of iBNA array with increasing linter from 50 to 150 nm. The solid tracing lines show the linear fit to the experimental data.

To construct a uniform iBNA with interparticle distance and density tunability, we employed the nano-sized building block self-assembly method using block-copolymer (BCP). In the self-assembly process, selective interactions between the specific chains of BCP and metal ions lead to the formation of nano-corona structures.^34^

The core of the corona comprises gold ions and hydrophilic parts of the BCP, and the exterior region of the corona is the hydrophobic part of the BCP extending into the hydrophobic solvent environment. This uniform geometric arrangement of the coronas at the nanoscale allows us to achieve a well-ordered corona array when the corona solution is deposited on the substrate. Next, in the process of BCP removal using the O2 plasma process, the Au ions are reduced, and finally, a gold nanostructured array is formed on the substrate. In this study, we employed polystyrene-*b*-poly (2-vinyl pyridine) (PS-*b*-P2VP) and a gold precursor to prepare the iBNA (**Experimental Sections**). The densities and the interparticle distance (*linter*) of iBNA were controlled by adjusting the molecular ratio between the BCP and Au (Figure 2B). Scanning electron microscopy (SEM) images of four different densities of the iBNA are shown in Figure 2C. In an iBNA area (*AiBNA*) of 10^4^ μm^2^, we observed a uniform distribution of Au nanoparticles on the substrate. At the lowest iBNA density (*ρPNA* = 50 particles/μm^2^), a two-dimensional (2d) hexagonal order of the iBNA on the substrate was observed. At a higher density of iBNA (*ρiBNA* = 800 particles/μm^2^), the orderness of the iBNA array decreased, but the particle size distribution remained constant (between 42 and 43.7 nm; Figure 2D).

Furthermore, the self-assembly of the BCP and metal ions enabled us to obtain a narrow size distribution of the iBNAs, regardless of the *ρiBNA* value changing from 50 to 800 particles/μm^2^. This narrow particle distribution resulted in a uniform interparticle distance (Figure 2E). The average interparticle distance decreased from 600 to ∼ 70 nm when *ρPNA* is increased from 50 to 800 particles/μm^2^, and the standard deviation (*σiBNA*) of particle distance is less than 7%.

When a plasmonic nanostructure is close to another (< 50 nm), an interparticle coupling between the two nanostructures causes a strong nanogap antenna effect.[34–37] The nanogap antenna effect results in a strong resonance peak shift compared to the isolated structure. As the two nanostructures come closer, the near field on one nanostructure can interact with that on the neighboring nanostructure. Thus, the overall electric (*E*) field originates from both the incident light field (*E0*) and the near field (*Enf*) of the neighboring nanoparticles. This nanogap antenna effect leads to high-intensity plasmonic coupling with an enhanced *E*-field but decreasing local resolution (Figure 2F). To evaluate the enhancement of the E-field, we calculated its distribution using finite element analysis (FEA). A high-intensity E-field is observed in the gaps between the iBNAs. Specifically, when the interspacing (*linter*) is 50 nm, the overlapping E-field results in an intensity five times higher than 600 nm. Conversely, when *linter* exceeds 300 nm, a distinct E-field appears on individual nanostructures.

Subsequently, we characterized the optical properties and sensitivity of the tunable iBNA. A short *linter* of the iBNA is associated with strong interparticle coupling effects, which enhance the *E*-field. This *E*-field enhancement around the iBNA was experimentally observed by dark-field microscopy on varying *linter* from 50 and 150 nm (Figure 2G). In contrast, the extinction peak of the low density (*linter* = 150 nm) iBNA occurred at *λmax* of ca. 550 nm, and the extinction peak shifted to *λmax* of 592 nm at a *linter* of ca. 50 nm. The full width at half maximum (FWHM) of the LSPR peak broadened from 165 to 190 nm as *linter* decreased from 150 to 50 nm. The larger variation of *linter* in the high-density iBNA mainly results in a broader FWHM.

Next, we quantified the sensitivity change as a function of *linter* by measuring the medium refractive index (*nm*) sensitivity as a function of *linter* from 50 to 150 nm (Figure 2H). We obtained the LSPR peak shift (*Δλ*) in refractive index units (RIU) and figures of merit (FOM = *Δλ/*(Full width at half maximum)). The LSPR peak shift linearly increased from 50 to 150 nm as *nm* increased from 1.33 to 1.47. In particular, we observed a maximum value of *Δλ* (50 nm) at a *linter* of 70 nm and *nm* of 1.47. The larger distribution of *linter* at the highest *ρPNA* induced a broader range of plasmonic coupling effects. The broader coupling led to broader spectral peaks and lower FOMs.

### 2.3. Highly sensitive and specific spatiotemporal imaging

While constructing iBNA, we first investigated sensing capabilities, including sensitivity, specificity, and spatial resolution (Figure 3). We confirmed the LSPR peak shifts from the plasmonic nanostructure array to the iBNA (aptamer conjugated plasmonic nanostructure array) and IL-6 on the iBNA (Figure 3A). We tested the aptamer conjugation effect on iBNA compared to the detection performance of controls such as only plasmonic nanostructure array, and control aptamer conjugated plasmonic nanostructure array (Figure 3B). In this test, after confirming spectral uniformity over the whole sample area (**Figure S1**), we loaded a simulated IL-6 sample onto the iBNA layer with *CIL-6* varying from 10^−1^ to 10^5^ pg mL^−1^. Only iBNA led to the detection intensity (*ΔI/I0*) increased as a function of *CIL-6*, while the others did not show visible changes. The iBNA assay achieved a limit of detection (LODiBNA) as small as 0.56 pg mL^−1^, whereas the gold standard Enzyme Linked Sandwich Assay (ELISA) exhibited an LODELISA of 27.6 pg mL^−1^ (**Supporting Information**). The ELISA test employed a commercial IL-6 antibody (Abcam, USA). Such a comparison shows that the iBNA has a superior concentration sensitivity (or LOD) for IL-6 over 50 times higher and 100 times rapid (**Figure S2**) than that of the ELISA assay.

**Figure 3.**
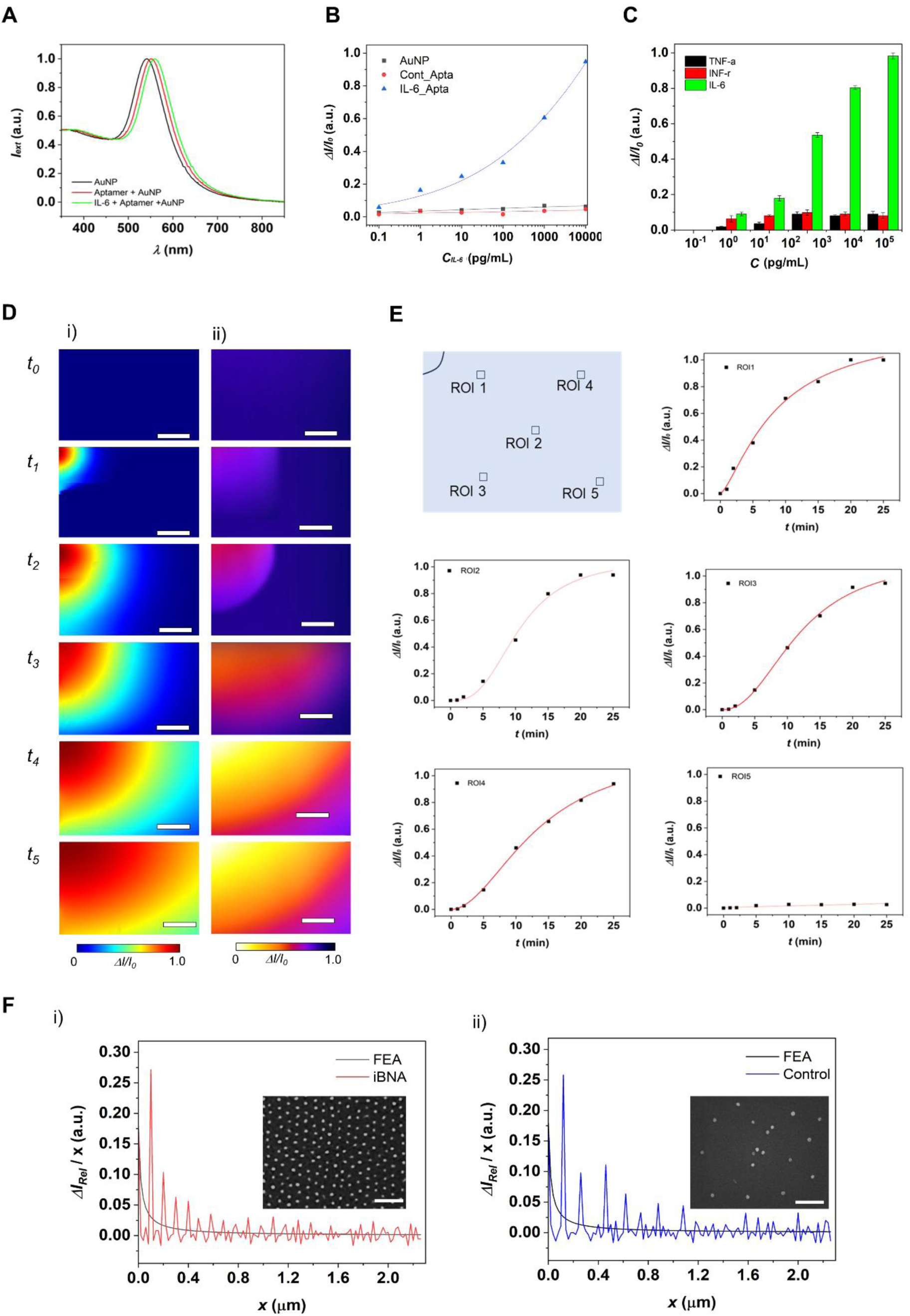
High spatiotemporal biomolecular imaging enabled by iBNA. A) LSPR peak of i) iBNA, ii) Ap-iBNA, iii) IL-6 loaded Ap-iBNA. B) Standard calibration curves (photocurrent change (ΔIph/Iph_0) versus CIL-6) of i) iBNA, ii) Control-Ap-iBNA, and iii) Ap-iBNA obtained using the LSPR spectroscopy setup. C) Signal (ΔI/I0) data of the iBNA obtained for a mixture of TNF-α, INF-γ, and IL-6 at various concentrations (n = 6). The concentrations for the background biomarkers: TNF-α, INF-γ, and IL-6 were varied so that their values were identical with that of IL-6 for each measurement. The data show high selectivity of iBNA to IL-6 is distinctly higher than those for the other biomarkers with a statistical significance (p ≤ 0.01). D) Spatiotemporal IL- 6 profiles acquired by i) numerical and ii) experimental approaches (scale bars = 100 μm) at t0 = 10 min, t1 = 30 min, t2 = 60 min, t3 = 90 min, t4 = 120 min, and t5= 150 min. The images show a gradual change in both intensity and area covered by IL-6 migration over selected time points. E) Dynamic signal change plot (ΔI/I0) versus CIL-6 at individual local points from ROI 1 to 5. F) IL- 6 intensity profile difference as a function of distance (ΔI/x), accompanied by inserted SEM images of (i) iBNA and (ii) the control substrate under identical concentration gradient conditions (scale bars = 1 μm).

Subsequently, we performed a control experiment to test the iBNA’s selectivity to the target biomarker, i.e., IL-6, in comparison with other different background biomarkers of INF-α and TNF-α with their concentrations ranging from 10^−1^ to 10^5^ ng mL^−1^ altogether (Figure 3C). In this test, we observed no noticeable changes in Δ*I/I0* for the concentrations of any biomarkers other than IL-6. This result indicates negligible cross-reactivity among these biomarkers, proving the capability of iBNA. Our method enables improved sensitivity compared to traditional fluorescence-based approaches.

We further demonstrated imaging of IL-6 profiles on iBNA (Figure 3D and **Figure S3**). When IL-6 was injected into a corner of an iBNA substrate, we acquired the profile images as a function of IL-6. As time passed, IL-6 diffusion occurred from the corner to the opposition direction, and the concentration change of IL-6 was spatiotemporally visualized. We also validated the diffusion of IL-6 using FEA (**Figure S4** and **Experimental Section**). The numerically acquired dynamic spatial profile of IL-6 is well-matched to the spatial profiles from iBNA. In addition, these spatiotemporal profiles allow dynamic binding curves at each region of interest (ROI) (Figure 3E). For example, ROI 1 is closer to the IL-6 injection location and shows more rapid IL- 6 saturation, whereas ROI 5 is farther away and shows slower IL-6 saturation.

We further estimated the spatial resolution of iBNA by plotting a differential intensity profile versus spatial distance, *ΔI/xiBNA* (Figure 3F **and Figure S5**). The estimated resolution from the average distance when *ΔI/xiBNA* is zero is ∼ 303 ± 5 nm. As a control, we also estimated the spatial resolution of an image via a plasmonic substrate with low density and uniformity. The control’s spatial resolution is ∼ 608 ± 150 nm at the same concentration gradient revealed by FEA result. Our observation here confirmed that the high density and uniform plasmonic nanostructure of the iBNA layer enables the sensitive LSPR spectral shift accompanying the variation of *CIL-6,* resulting in high spatiotemporal resolution imaging. Furthermore, we have verified that iBNA effectively captures the temporal profiles of IL-6 across varying injection directions (**Figure S6**).

### 2.4. Spatiotemporal imaging of cytokines secreted from a single cell

We constructed iBNA and demonstrated spatiotemporal imaging of cytokine secretion from single cells (Figure 4). Jurkat T cells, a widely used human leukemia cell line for studying T-cell receptor signaling pathways, were employed as a model to demonstrate cytokine secretion imaging.[38] We used phorbol 12-myristate 13-acetate (PMA) and ionomycin to stimulate Jurkat T cells to produce stimulation responses independent of the T cell receptor (TCR).[12,16] Then, stimulated single Jurkat T cells were loaded into the nanoplasmonic microarray device for dynamic cytokine imaging (Figure 4A, **Figure S7**, and **Experimental Section**). Figure 4B shows acquired dynamics cytokine profile images from a stimulated single Jurkat T cell and a non-stimulated one. After two hours, the IL-6 profile is shown as secreted from the stimulated Jurkat T cell, while the unstimulated Jurkat T cell leads to invisible profiles of IL-6 over time. These images indicated that the iBNA enables them to visualize the dynamic profile of IL-6 from the same single cell. To validate the high spatial resolution of the IL-6 profile obtained via iBNA, we conducted a comparative analysis of IL-6 profiles generated using iBNA and those derived from sparsely populated and non-uniform plasmonic nanostructure arrays (Figure 4C). The iBNA demonstrated its capability to resolve the local distribution of IL-6 secreted by a stimulated Jurkat T cell. In contrast, a low-density yet uniform plasmonic array yielded a non-smooth profile with diminished resolution, whereas low-density, non-uniform plasmonic arrays produced irregular and inconsistent imaging results.

**Figure 4.**
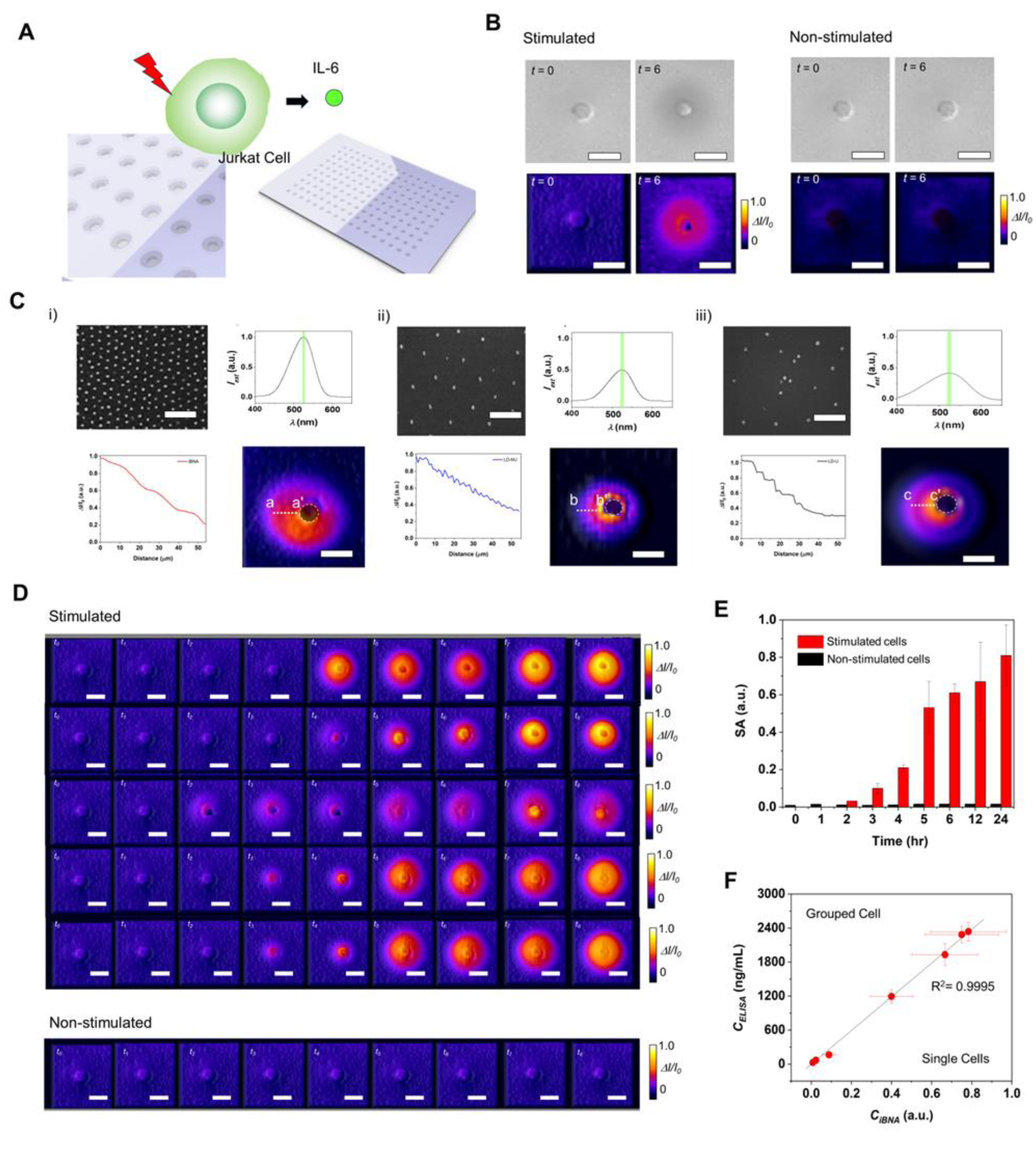
High spatiotemporal imaging of proteins secretion of single Jurkat T cells. A) Schematic of high spatiotemporal imaging cytokine secretion of stimulated single Jurkat T cell. B) Bright-field optical (top) and iBNA (bottom) images of single Jurkat T cells at t = 0 and t = 6 hours, respectively, showing i) stimulated and ii) non-stimulated Jurkat T cells. Microwell arrays are used to track the cells’ positions (scale bars = 10 μm). C) SEM images (scale bars = 500 nm) and LSPR spectra of (i) iBNA, (ii) uniform low-density plasmonic arrays, and (iii) non-uniform low-density plasmonic arrays, along with intensity profiles corresponding to the a-a’, b-b’, and c-c’ lines in the IL-6 profiles obtained from each nanostructure array. D) Dynamic IL-6 secretion profiles of five representative stimulated Jurkat T cells (top) and a non-stimulated Jurkat T cell (bottom) showed dynamic IL-6 secretion profiles over a full day (scale bars = 20 μm). Heterogeneous changes in secretion over specific time points are indicated by these. E) Dynamic profiling of IL-6 secretion from individual stimulated and non-stimulated Jurkat T cells (n=100) assessed through surface area (SA) analysis. F) Linear regression correlating the IL-6 SA results from iBNA imaging for individual stimulated Jurkat T cells (n=100) with the ELISA data from grouped cells in petri dishes (cell density ∼10⁶ cells/mL). Measurements were made at t = 0, 1, 2, 3, 4, 5, and 6 hours, and the IL-6 concentrations varied from 10⁻² to 2,400 ng/mL

In addition, we confirmed heterogeneous characteristics of the IL-6 secretion from different Jurkat T cells (Figure 4D). The initial section of IL-6 from each cell occurs around two or three hours; however, the simulated dynamic cytokine profiles from five distinct single Jurkat-T cells reveal diverse locations and time-sequence patterns. The iBNA permits cytokine profiles from single cells, allowing for the acquisition of secretion heterogeneity amongst singles.

The high-throughput imaging and analytical capabilities of iBNA enable the detailed characterization of secretion profiles at the single-cell level and provide valuable insights into dynamic secretion patterns and overarching trends of multiple cells. By quantifying the area of IL- 6 profiles from the acquired images, we constructed a cytokine intensity plot as a function time from numerous single cells (Figure 4E). The macro trends were derived based on the individual cell-level patterns of IL-6 secretion observed in **Figure S8**. For the stimulated single Jurkat-T cells, as we observed from the images, the cytokine intensity increases around 2hr or 3hr. The secretion continuously increases to 12 hr and saturates between 12 and 24 hrs. However, the cytokine intensity of non-stimulated single Jurkat-T cells results in invisible change over time.

Furthermore, we validated the cytokine secretion trends obtained with a gold standard approach, ELISA. We note that iBNA allows us to use the same single cell over time, while ELISA needs multiple cell samples, enabling the collection of medium samples each time. Dynamic analysis of IL-6 for both iBNA and ELISA was performed, and we constructed a regression to correlate the results from iBNA and commercial ELISA assays (Figure 4F). The correlation shows a strong linear relationship (R^2^ = 0.9938). This relationship indicates that the trends obtained in IL-6 on iBNA have been validated well with the gold standard approach over time.

### 2.5. Study cell-to-cell communication by imaging cytokine release

By utilizing iBNA to analyze the dynamic cytokine profile of a single Jurkat T cell, we conducted imaging that reveals how cytokine secretion mediates communication between individual cells (Figure 5). We selected the Jurkat T and CD4+ T cells from which CD4+ T cells can be differentiated as a model to study the mechanisms of cell-cell communication (Figure 5A and **Figure S9**).[6,39] IL-6 plays a critical role in the differentiation and function of CD4+ T cells by binding to the IL-6 receptor on their surface and activating intracellular signaling pathways that promote their development and activity. In this study, the process of CD4+ T cells receiving IL-6 involves four steps. IL-6 is produced by the Jurkat T cell in response to PMA stimulation. CD4+ T cells express the IL-6 receptor (IL-6R) on their surface. When IL-6 binds to IL-6R, it triggers a signaling cascade within the cells.[39–42] The binding of IL-6 to IL-6R activates intracellular signaling pathways, including the Janus Kinase (JAK)/ Signal Transducer and Activator of Transcription (STAT) pathway. This leads to the phosphorylation and activation of STAT3, a transcription factor1. Activated STAT3 trans locates to the nucleus and promotes gene expression in CD4+ T cell differentiation and function. This includes the production of other cytokines like IL-17, which further amplifies the inflammatory response (Figure 5B).

**Figure 5.**
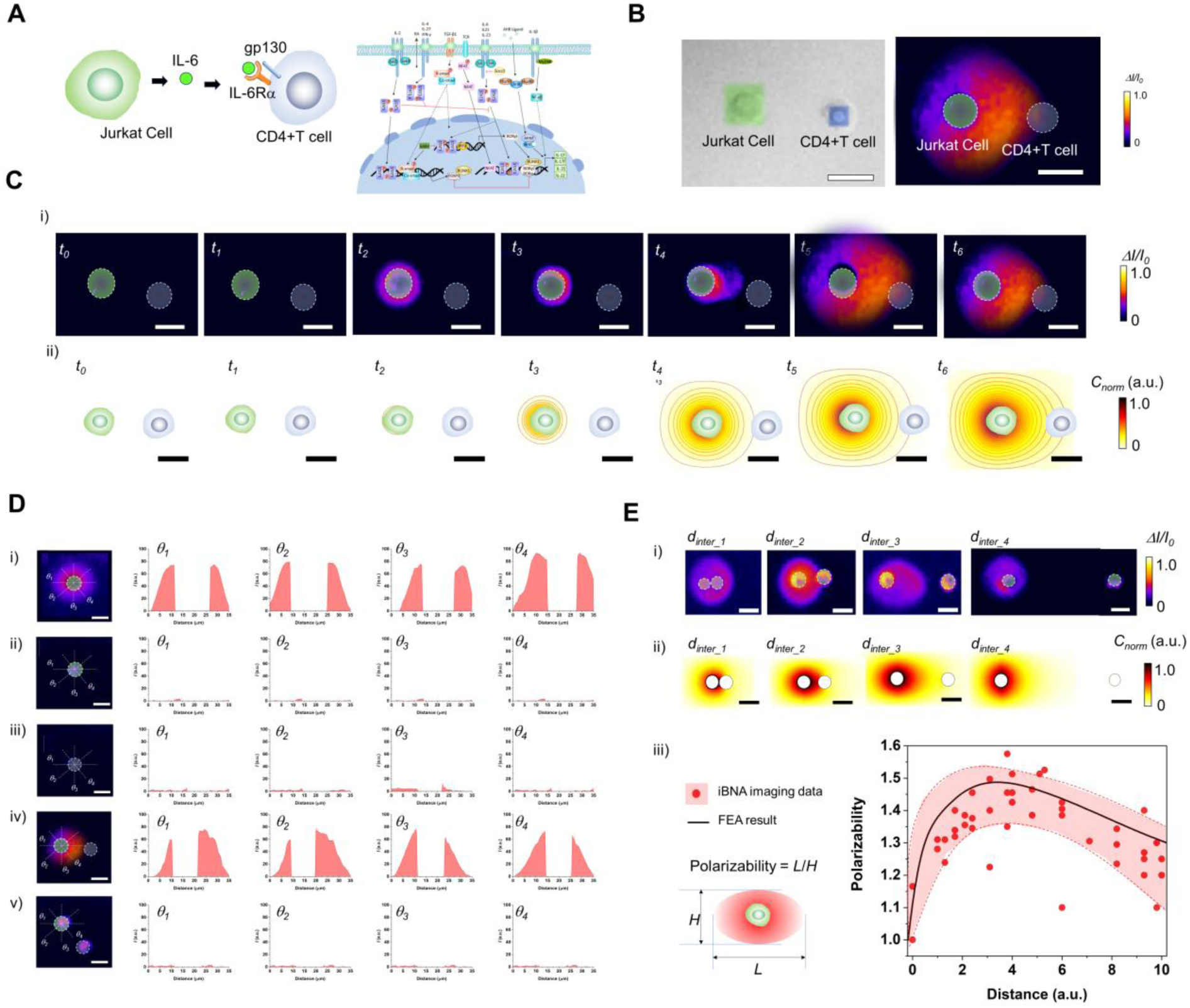
Spatiotemporal Imaging of Cell-Cell Communication via Cytokine Secretion. A) Schematic representation of cytokine-mediated cell-cell communication between a stimulated Jurkat T cell and a CD4+ T cell. B) Visualization and confirmation of the Jurkat T cell and CD4+ T cell via (i) brightfield-overlapped fluorescence images and (ii) iBNA imaging using an integrated microscope setup with fluorescence imaging (scale bars = 20 μm). To differentiate cell types, Jurkat T cells and CD4+ T cells were labeled with FITC and DAPI, respectively. C) Dynamic IL- 6 secretion profiles between a stimulated Jurkat T cell and a CD4+ T cell, obtained through imaging and numerical analysis (scale bars = 20 μm). In the numerical model, IL-6 secretion from the Jurkat T cell and receptor distribution on the CD4+ T cell membrane were assumed to be uniform. D) IL-6 distribution images and concentrations along the central axis of single cells: (i) stimulated Jurkat T cell, (ii) non-stimulated Jurkat T cell, (iii) CD4+ T cell, (iv) stimulated Jurkat T cell with a CD4+ T cell, and (v) non-stimulated Jurkat T cell with a CD4+ T cell, analyzed at angles of 0°, 45°, 90°, and 135° relative to the central axis (scale bars = 20 μm). E) i) Experimentally and ii) numerically determined IL-6 profiles as a function of intercellular distance between stimulated Jurkat T cells and CD4+ T cells (scale bars = 20 μm). iii) IL-6 polarization profiles as a function of intercellular distance between stimulated Jurkat T cells and CD4+ T cells. Data points (red spheres) represent single-cell measurements, while the black curve reflects numerically derived profiles.

Figure 5C shows dynamic images of IL-6 profiles secreted from a Jurkat T cell stimulated to CD4+ T cell. IL-6 intensity rises around the Jurkat T cell after two hours, and the IL-6 distribution symmetrically widens over time. The distribution of IL-6 toward CD4+ T cells starts at 5 hours and gradually becomes polarized (Figure 5C). In addition, we validated such polarized IL-6 migration patterns from the stimulated Jurkat T cell and CD4+ T cell using FEA (Figure 5D and **Figure S10**). A uniform distribution appears when IL-6 secretion begins. The distribution indicates that the polarized profile rises over time as more IL-6 is secreted and approaches the CD4+ T cell. The FEA results are consistent with the dynamic patterns of IL-6 between Jurkat T cells and CD4+ T cells that were experimentally acquired on iBNA.

Figure 5E shows plots of IL-6 level, a cross with 0, 45, and 90° angles at the center of cells, including a stimulated Jurkat T cell, a non-stimulated Jurkat T cell, CD4+ T, a stimulated Jurkat T cell with a CD4+ T cell, and a non-stimulated Jurkat T cell with a CD4+ T cell. The case of a stimulated Jurkat T cell and a CD4+ T cell only shows the higher and polarized level of IL-6 toward the CD4+ T cell. In contrast, a stimulated Jurkat T-cell results in a symmetric IL-6 pattern; other cases do not show any profiles. The polarized IL-6 profile highlights the role of IL-6 as a mediator in the communication between a stimulated Jurkat T cell and a CD4+ T cell. IL-6 is a cytokine that plays a pivotal role in immune responses, particularly in the differentiation and function of CD4+ T cells. We further characterized the polarized pattern IL-6 during cell-to-cell communication as a function of intercellular distance (Figure 5F**, G**). As a stimulated Jurkat T cell is closer to the CD4+ T cell, the IL-6 profiles reveal symmetric patterns, while the longer distance between the two cells leads to polarized patterns. However, the polarized pattern is decreased when the intercellular distance is longer than the critical distance. From the comprehensive analysis of iBNA of multiple cell-to-cell communication cases, we observed polarized IL-6 profiles between the stimulated Jurkat T cells and CD4+ T cells. This polarized IL- 6 trend as a function of intercellular distance also matches that obtained from FEA results. The polarized IL-6 profile suggests that IL-6 production is concentrated in specific contexts or interactions (**Figure S11**). Finally, iBNA allowed us to acquire direct and dynamic images of the polarized cytokine profiles during cell-cell communications. This polarization could enhance the specificity and efficiency of communication between Jurkat T cells and CD4+ T cells, potentially influencing immune responses or inflammatory processes.[43,44]

## 3. Conclusion

In conclusion, we created iBNA using a self-assembled gold nanostructure array functionalized with aptamer receptors, enabling 100 times more sensitive, significantly specific, and prompt IL- 6 detection than traditional methods. The imaging principle involves tuning the delivery of incident light through the iBNA, leveraging biomolecular surface binding-induced localized plasmonic resonance shifts. This tuning depends on the secreted cytokine concentration on the iBNA, which was detected optoelectronically. With the developed iBNA, we achieved real-time spatial resolution imaging of secreted IL-6 from live single cells. Additionally, we successfully visualized dynamic cell-cell communication by imaging the IL-6 secretion profile over time. Our bioinspired nanoplasmonic platform represents a transformative advancement in real-time cellular secretion analysis. The high spatiotemporal resolution and label-free nature of iBNA make it a promising tool for applications ranging from immunological research to precision medicine. Future research should explore its adaptability for multiplexed protein detection and clinical validation in disease monitoring. Additionally, integrating machine learning algorithms could enhance signal interpretation, enabling automated and predictive analysis for broader biomedical applications.

## 4. Experimental Sections

*Chemicals:* We purchased gold (III) chloride trihydrate, toluene, isopropanol, C-10, BSA, TCEP, MCH, K₃[Fe(CN)₆], K₄[Fe(CN)₆], PBS buffer, and MgCl₂ from Sigma Aldrich. We obtained polystyrene-b-poly (2-vinyl pyridine) from Polymer Science, Anti-Interleukin 6 (IL-6) Aptamer (CTApt-217) with 5’ (Thiol C6 S-S) group from Creative Biolabs, cytokines from Thermo Fisher Scientific, and PDMS along with its curing agent from Corning. Our laboratory produced nano pure water in-house (Details are described in **Supporting Information.**).

*iBNA Fabrication:* We fabricated iBNA using block copolymer nanolithography. We dissolved polystyrene-b-poly(2-vinylpyridine) in toluene, allowed it to self-assemble with gold chloride trihydrate, and formed micelles. We spin-coated the micelles onto piranha-cleaned glass slides and treated the coated slides with O₂ plasma. SEM analysis confirmed the resulting morphology (Details are described in **Supporting Information.**).

*Bio-Conjugation Process:* We prepared aptamers by heating, cooling, and incubating them with TCEP based on a standard protocol. We applied the treated aptamers to Au NP arrays, facilitated thiol-gold binding through incubation, and treated the arrays with MCH to reduce nonspecific adsorption (Details are described in **Supporting Information.**).

*Characterization of LSPR Biosensors*: We mounted the LSPR biosensor chip on a motorized stage to align sensing spots and automate scanning. A dark-field condenser collected spectra from iBNA, which we captured using a spectrometer (Details are described in **Supporting Information.**).

*Finite element analysis:* In electromagnetic field calculation, we simulated electromagnetic fields around iBNA using COMSOL Multiphysics. We incorporated the material’s permittivity and polarization and based the nanostructure features on SEM images. In FEA modeling of cytokine diffusion, we simulated the movement of cytokines (small, diffusible proteins) through the medium, considering factors like geometric arrangement and diffusion to understand how they influence immune cell communication and responses (**Details are described in Supporting Information**).

*Imaging Setup:* We used an inverted microscope equipped with a CMOS camera for near-infrared imaging. Nikon software controls automated scanning for efficient image acquisition (Details are described in **the supporting information**).

*Single-Cell Loading:* We fabricated PDMS microwells, cleaned them, and prefilled them with glycerol-supplemented media. Using cellenONE X1 technology, we dispensed centrifuged and adjusted cells into microwells with precise control (Details are described in **Supporting Information.**).

*ELISA:* We quantified IL-6 using conventional ELISA. We performed sequential antibody incubations, applied TMB substrate, and measured absorbance at 450 nm to determine cytokine levels (Details are described in **Supporting Information.**).

*Cellular Model Preparation:* We cultured Jurkat cells with PMA and Ionomycin to stimulate cytokine secretion. After incubation, we collected supernatants with minimal sampling and applied them to the LSPR chip for analysis (Details are described in **Supporting Information.**).

## Acknowledgements

This work was supported by the National Institute of Health, NIH-NIGMS program (151528), and the Academic Research Fund at the University of Michigan.

