## Supporting Information for "High-Spatiotemporal Imaging of Protein Secretion During Cell-to-Cell Communication via Integrative Biosensing Nanoplasmonic Array"

**List of contents**

**1. Experimental Section**

**2. Supporting Figures**

### 1. Experimental Section

**Chemicals:** Gold(III) chloride trihydrate, toluene, isopropanol, 10-carboxy-1-decanethiol (C-10), bovine serum albumin (BSA), Tris(2-carboxyethyl)phosphine (TCEP), 6-mercaptohexanol (MCH), potassium ferricyanide ($K_3[Fe(CN)_6]$), potassium ferrocyanide ($K_4[Fe(CN)_6]$), phosphate-buffered saline (PBS) buffer, magnesium chloride ($MgCl_2$), gold(III) chloride trihydrate, and toluene were sourced from Sigma Aldrich. Polystyrene-b-poly(2-vinylpyridine) (S: 213,000; VP: 153,000) was obtained from Polymer Science. Anti-Interleukin 6 (IL-6) Aptamer (CTApt-217) wih 5' (Thiol C6 S-S) group was supplied by Creative Biolabs, while IL-6, TNF-α, and IFN-γ were procured from Thermo Fisher Scientific. Polydimethylsiloxane (PDMS) elastomer and its curing agent were purchased from Corning. In-house production provided nanopure deionized water (resistance: 18.1 MΩ·cm).

**iBNA fabrication:** A uniform, high-density array of gold plasmonic nanostructures (iBNA) was fabricated through block copolymer nanolithography, as detailed in previous studies.33-34 Polystyrene-b-poly(2-vinylpyridine) (S: 213,000 units; VP: 153,000 units) was dissolved in toluene at a concentration of 4 mg/mL and stirred overnight. Gold(III) chloride trihydrate powder, with a molar ratio of 0.4 per vinyl pyridine unit, was subsequently added and stirred for 72 hours to enable self-assembly within the cores of polystyrene-b-poly(2-vinylpyridine) micelles. The micelle solution was then spin-coated onto glass slides pre-cleaned with piranha solution, forming a uniform monolayer. Following the coating process, the glass slides underwent treatment with oxygen plasma (500 W, 0.3 mbar) for 30 minutes. Finally, the morphology of the iBNA was confirmed through scanning electron microscopy (SEM).

**Bio-conjugation process:** To prepare the aptamer solution, it was first diluted to a concentration of 100 μM using ultrapure water. Aptamer uncoiling was then achieved by heating the DNA solution to 95 °C for 12 minutes, followed by rapid cooling in ice-chilled water for 30 minutes. Subsequently, the DNA aptamer sequence was incubated with TCEP for 2 hours at room temperature and further diluted to various concentrations using ultrapure water. A 5 μL aliquot of the pretreated aptamer solution was then applied to small sections of pre-fabricated Au NP arrays under wet conditions. The substrate was incubated at 4 °C for 10 hours to facilitate thiol-gold binding. After incubation, the substrate was rinsed to remove unbound DNA and immersed in a $10^{-7}$ M MCH blocking agent for 2 hours to prevent nonspecific adsorption on the Au NP surface. Finally, the substrate was rinsed again and dried under a flow of argon.

**Characterization of the optical properties:** The fabricated LSPR biosensor microarray chip was mounted on a motorized stage (ProScan, Prior Scientific) to facilitate precise positioning of the on-chip sensing spot and enable automated signal scanning. A dark-field condenser (NA = 1.45, MBL12000, Nikon) was positioned in close proximity to the backside of the glass substrate using immersion oil. Extincted light from the iBNA was collected using a 20× objective lens located beneath the chip. The corresponding spectra were acquired using a spectrometer (Ocean Optics, USB 4000).

**Calculation of the electric field:** We simulated the electromagnetic fields surrounding the iBNA structure using finite element analysis (FEA) in COMSOL Multiphysics by solving the Helmholtz wave equation. To account for its rounded geometry, hybrid mesh structures were designed for the iBNA. The relative permeability ($\mu_r$ = 1) and complex permittivity ($\varepsilon_r$ = f(λ)) of gold were incorporated into the simulations. A polarization vector was aligned parallel to the iBNA, while the wave vector (k-vector) was oriented perpendicularly to the plane of the iBNA structure. To minimize reflections, perfect absorption was assumed at the outer boundary by implementing a perfectly matched layer (PML) and an integration layer within concentric space. The diameter of the iBNA ($d_{iBNA} \approx$ 42 nm) was selected based on the SEM images presented in Figure 2.

**Imaging setup**: An inverted microscope (Olympus IX73) served as the primary optical platform for label-free cell secretion analysis and was equipped with a customized microscope cell incubator (Life Imaging Services). For plasmonic intensity imaging, a collimated near-infrared LED (Thorlabs, M850L3-C5) controlled by an LED driver (Thorlabs, LEDD1B) provided narrowband illumination. Considering the long-term and time-laps imaging, ultra-low noise level from a camera is critical. To ensure steady performance overtime, the images from the probes were acquired at different positions from a CCD camera (Pixis). The microscope stage was operated via an in-house code, facilitating automated scanning of multiple FOVs for high-throughput imaging.

**Single cell loading in the detection device:** Microwell structures were prepared by attaching a PDMS micromesh to the gold nanohole array chips for single cell seeding. The PDMS structures were fabricated using standard photolithography and soft lithography techniques. Each microwell measured 200 μm in diameter and 50 μm in height, providing a unit volume of 1.5 nL. The PDMS device was cleaned through sonication in 70% ethanol and dried with pressurized nitrogen prior to cell seeding. Single cells were isolated and dispensed into the microwells using the advanced cellenONE X1 technology (SCIENION), which combines piezoelectric liquid dispensing with sophisticated image processing for deterministic single-cell deposition. To prevent evaporation, each microwell was prefilled with cell culture medium containing 1% v/v glycerol. Cell suspensions were centrifuged at 410 × g for 5 minutes and washed twice with serum-free media to remove secreted materials. The cell density was adjusted to 2–3 × $10^5$ cells per mL, and 50 μL of the suspension was transferred to a 384-well plate, compatible with the dispenser. Using a piezoelectric voltage of 65 V and a pulse duration of 48 μs, 300 pL droplets containing cell culture media were formed. Finally, 10 μL of cell suspension was loaded into the dispensing nozzle, enabling highly precise single cell seeding into microwells.

**Enzyme-linked sandwich assay (ELISA):** Enzyme-linked immunosorbent assay (ELISA) serves as the gold standard for quantifying IL-6 in cellular media. A cell culture medium of 50 μL containing Jurkat T cells at a concentration of 2.5 × $10^6$ cells/mL was collected hourly and directly loaded onto the ELISA assay plate for cytokine quantification, utilizing a conventional ELISA kit (Abcam, Cambridge, MA, USA). The ELISA plates were pre-blocked with blocking buffer (Thermo Scientific, Rockford, IL, USA) at room temperature for 2 hours. Serum was treated with DNase and subsequently added to the wells alongside 80 μL of blocking buffer, followed by a 2-

hour incubation at room temperature. After four washes, an anti-IL-6 polyclonal antibody (Abcam, Cambridge, MA, USA) was introduced as the detecting antibody and incubated at room temperature for 2 hours. This was followed by four additional washes and the addition of an anti-rabbit peroxidase-labeled secondary antibody, which was incubated in the wells at room temperature for 1 hour. Unbound secondary antibodies were removed via four thorough washes. The plates were treated with 3,3',5,5'-Tetramethylbenzidine (TMB) substrate in the dark for 20 minutes, after which a stop solution (R&D Systems Inc., Minneapolis, MN, USA) was added. The levels of IL-6 were quantified by measuring absorbance at $\lambda = 450$ nm.

**Preparation of cellular model:** A cell culture medium (2 mL) containing Jurkat cells at a concentration of $2.5 \times 10^6$ cells/mL was transferred into a well of a 6-well plate. To stimulate cytokine secretion, a mixture of PMA (100 ng/mL, Sigma-Aldrich) and Ionomycin (1000 ng/mL, Sigma-Aldrich), dissolved in deionized water, was added to the prepared cell suspension. The cells were then incubated for 2 hours. A 10 μL aliquot of supernatant was collected from the cell culture medium within the 6-well plate. This collected volume, constituting less than 1% of the total cell culture medium, minimized changes in cytokine concentration during sampling. Of the collected supernatant, 6 μL was directly loaded onto the LSPR biosensor microarray chip for cytokine quantification.

### 2. Supporting Figures and Captions

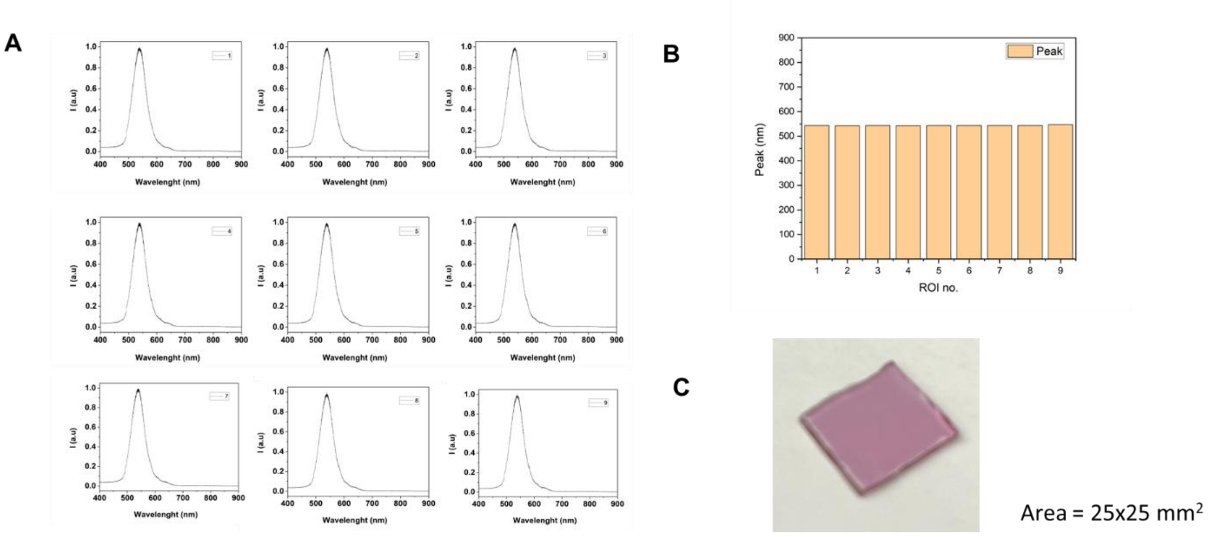


**Figure S1. Spectral uniformity of the iBNA sample. A)** Extinction spectra measured from nine different regions of interest (ROIs) on the iBNA sample. **B)** Peak positions of the spectra corresponding to the nine ROIs. **C)** Photograph of the iBNA sample (dimensions: length × width = 25 × 25 $mm^2$).

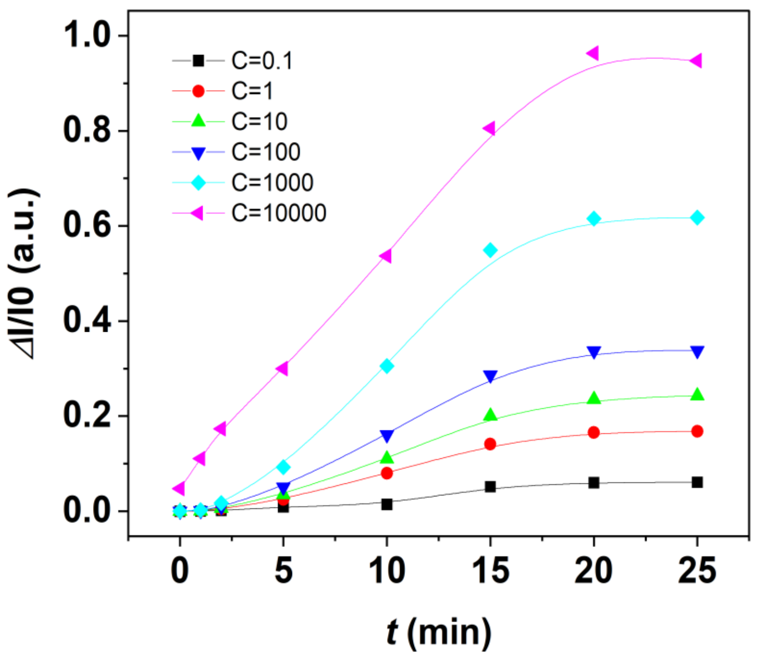


**Figure S2**. Time evolution of the normalized relative intensity ΔI/I0 signal for various levels of IL-6 concentration (unit: ng/mL), where I0 is the intensity value at C = 0. Each signal curve is measured during an assay incubation process over time.

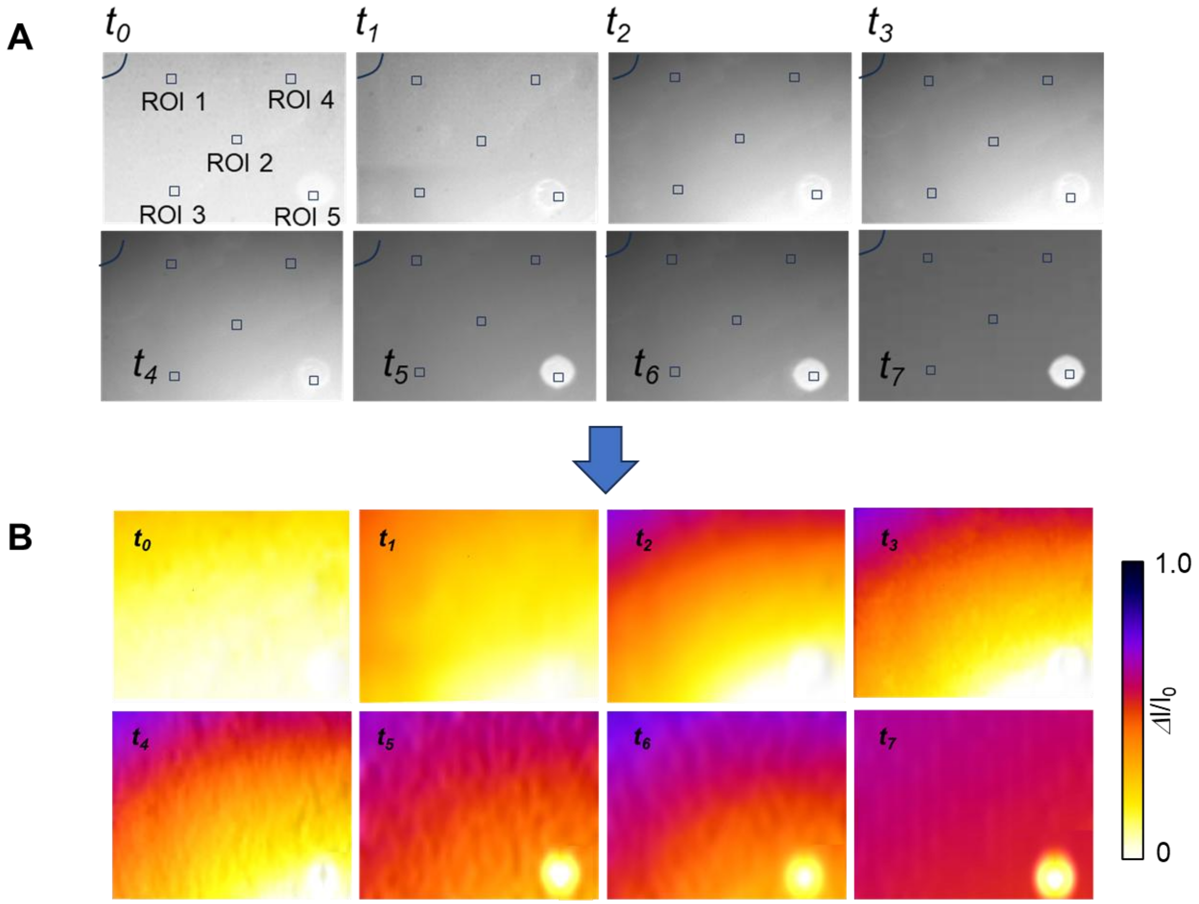


**Figure S3**. Monochrome and converted IL-6 profiles after injecting IL-6 at the top-left corner at $t_0$ = 10 min, $t_1$ = 30 min, $t_2$ = 60 min, $t_3$ = 90 min, $t_4$ = 120 min, and $t_5$= 150 min.

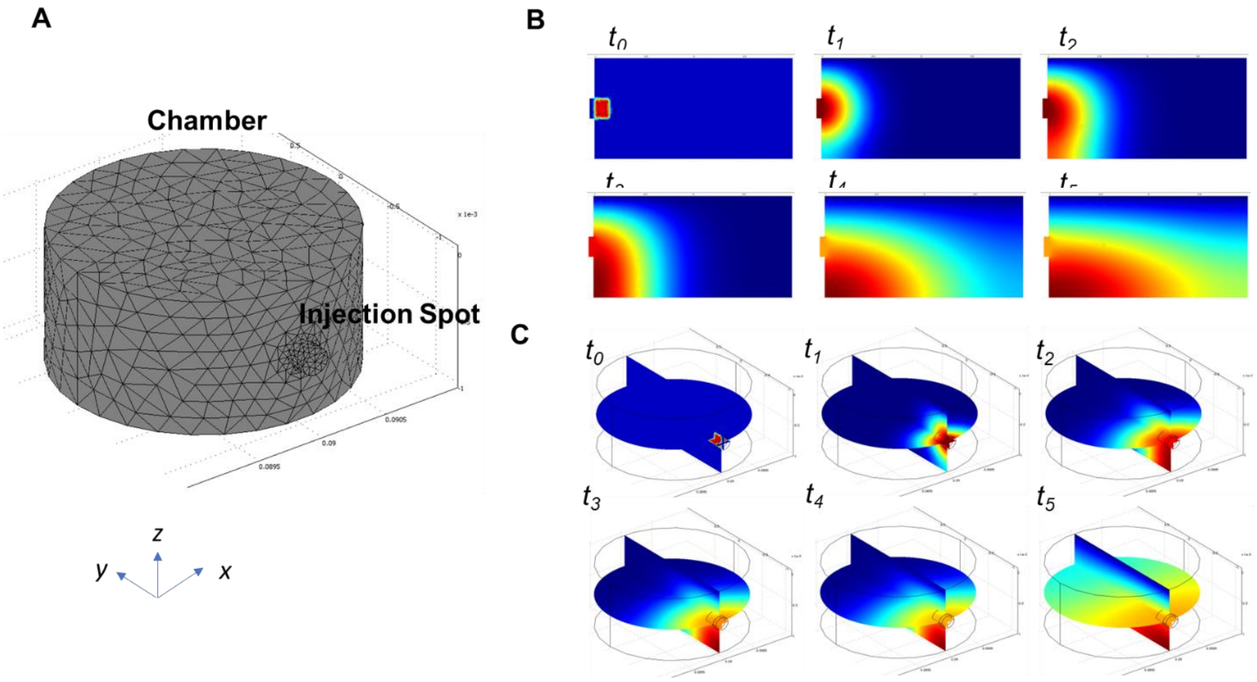


**Figure S4**. Finite element analysis of IL-6 diffusion model show. A) Constructed computational mesh structure. Obtained IL-6 concentration profile from B) side and C) perspective views at $t_0$ = 10 min, $t_1$ = 30 min, $t_2$ = 60 min, $t_3$ = 90 min, $t_4$ = 120 min, and $t_5$= 150 min.

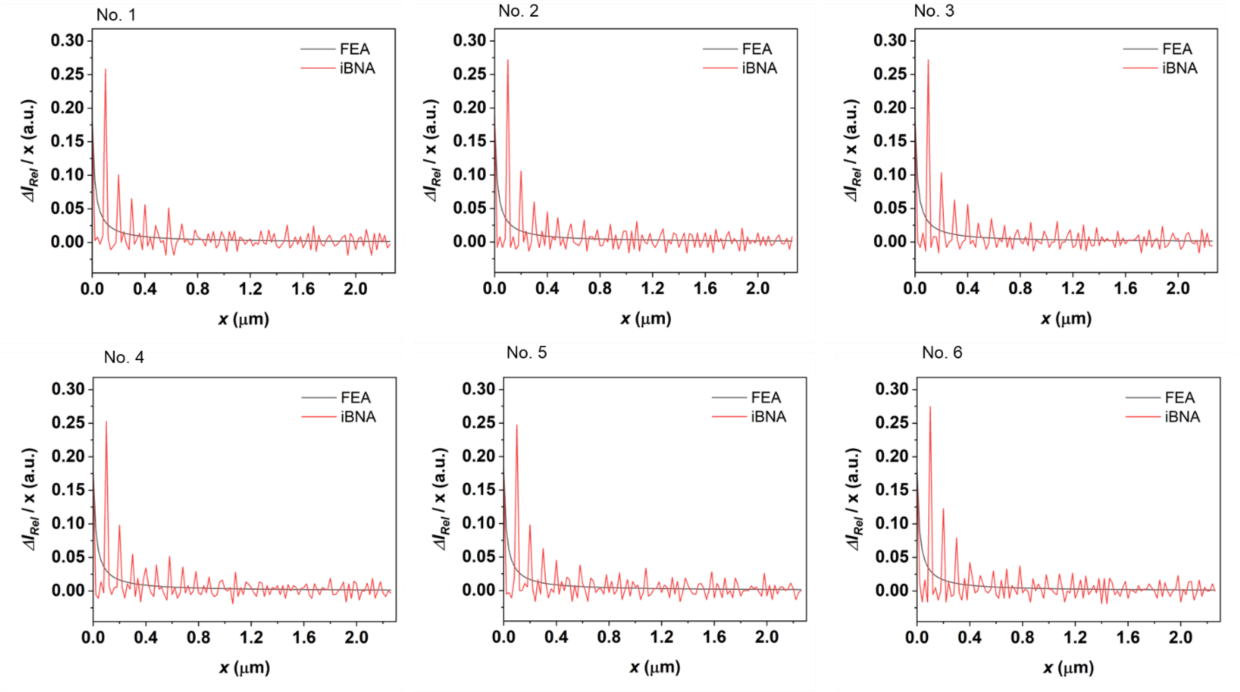


**Figure S5**. IL-6 intensity profile difference as a function of distance (*ΔI*/*x*) under identical concentration gradient conditions at six different locations.

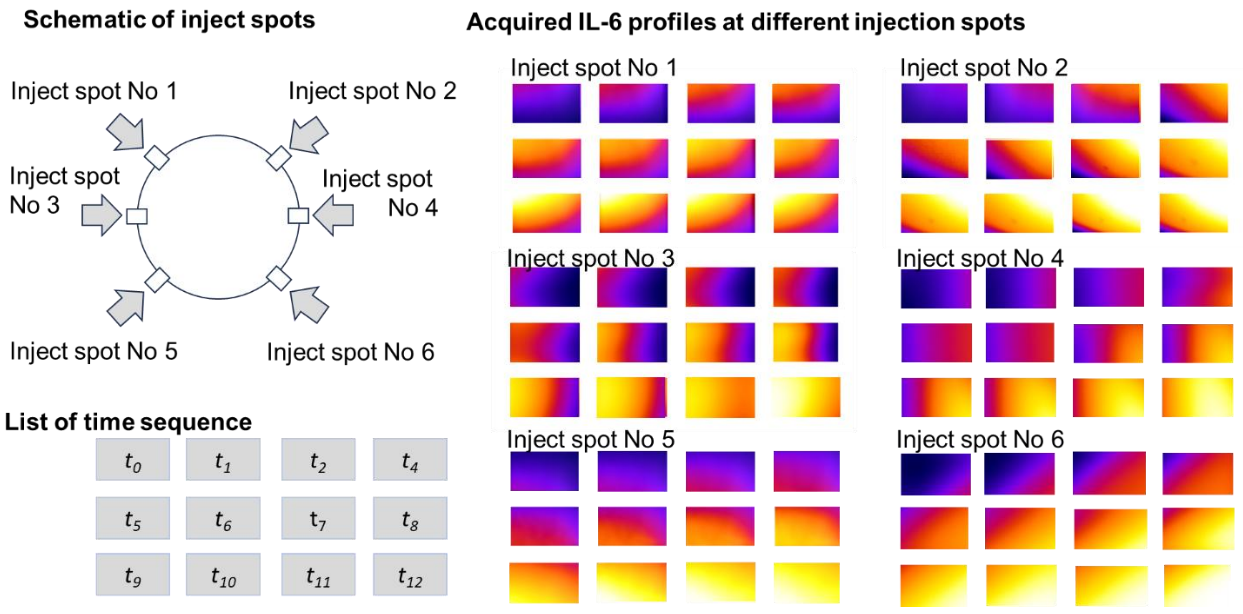


**Figure S6**. Acquired IL-6 profiles after injecting IL-6 at six different spots from i) to vi), respectively at $t_0$ = 10 min, $t_1$ = 30 min, $t_2$ = 60 min, $t_3$ = 90 min, $t_4$ = 120 min, $t_5$= 150 min, $t_6$= 180 min, $t_7$= 210 min, $t_8$= 240 min, and $t_9$= 300 min.

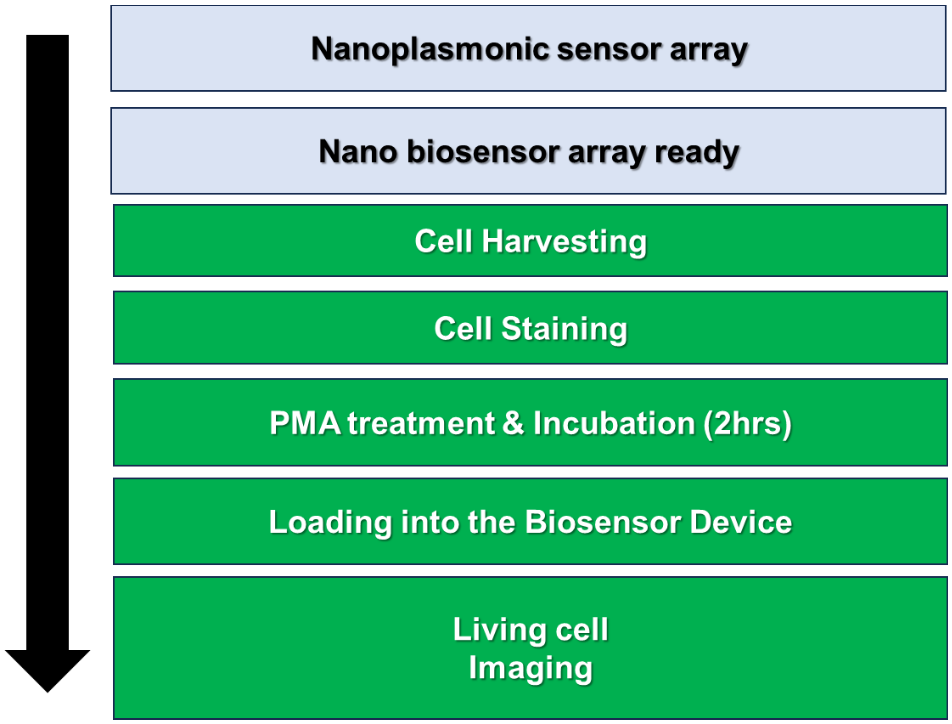


**Figure S7**. Flow chart describing experimental procedure from preparation of Nanoplasmonic sensor array to imaging.

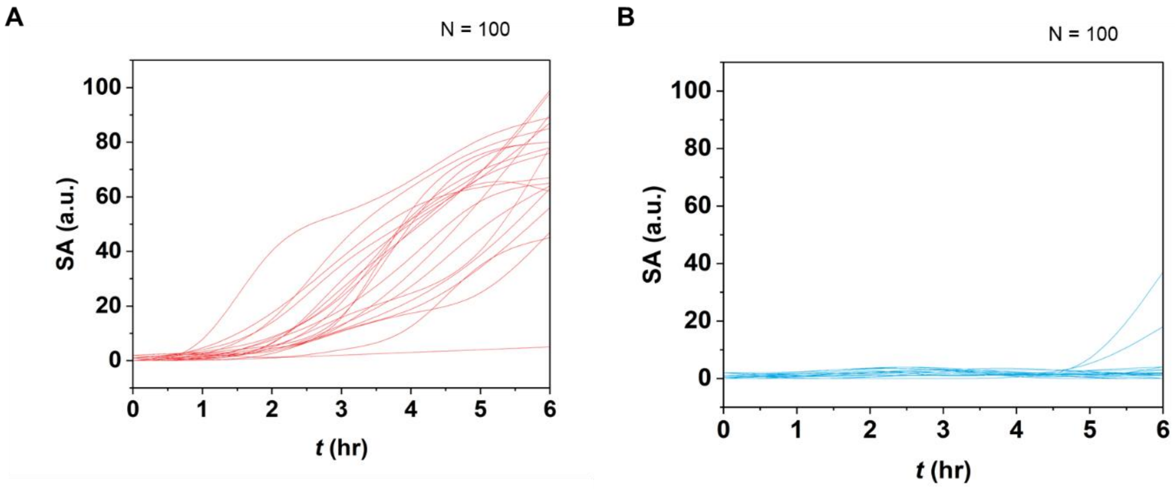


**Figure S8.** IL-6 secretion SA quantified from individual A) stimulated and B) non-stimulated Jurkat T cells (n=100) as a function of time.

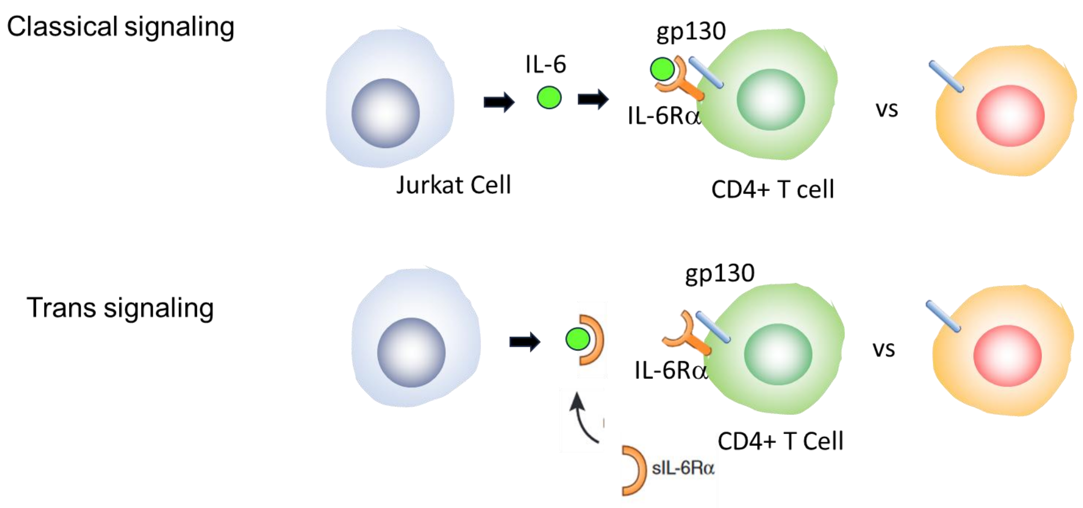


**Figure S9**. Schematic of cell-to-cell communication models based on classical and trans signaling.

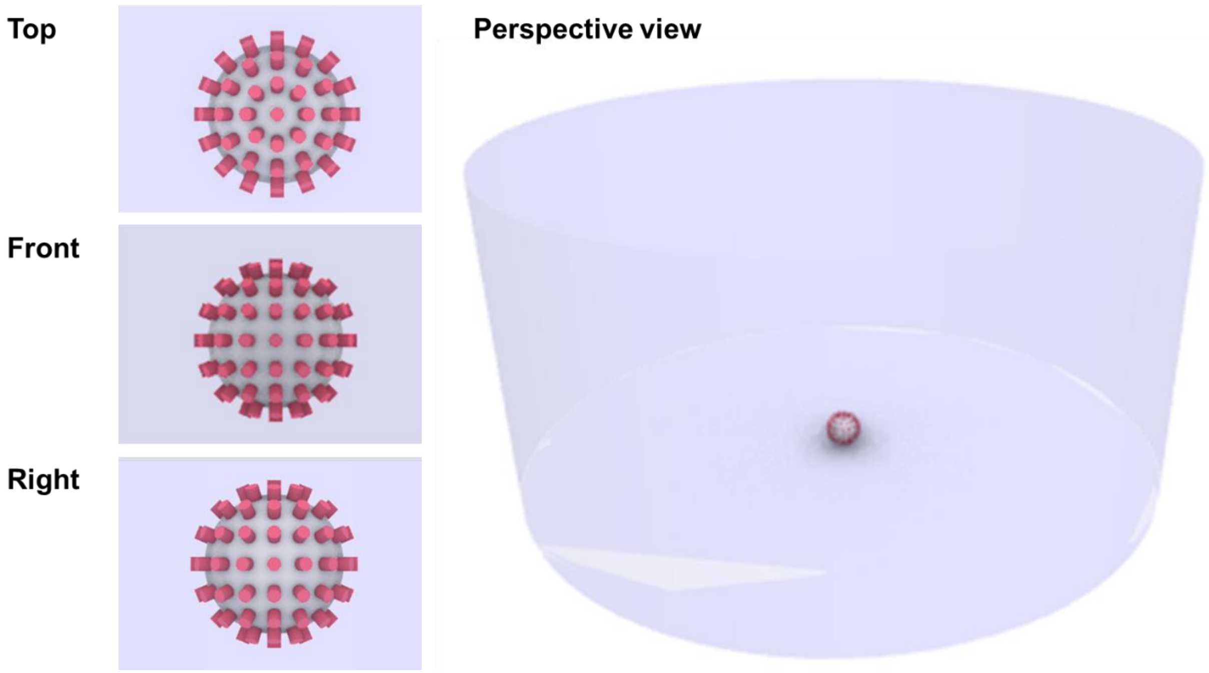


**Figure S10.** Schematic image of a single cell geometry that is applied in numerical analysis. In the model, 52 secretion spots uniformly distributed on the cell surface were applied.

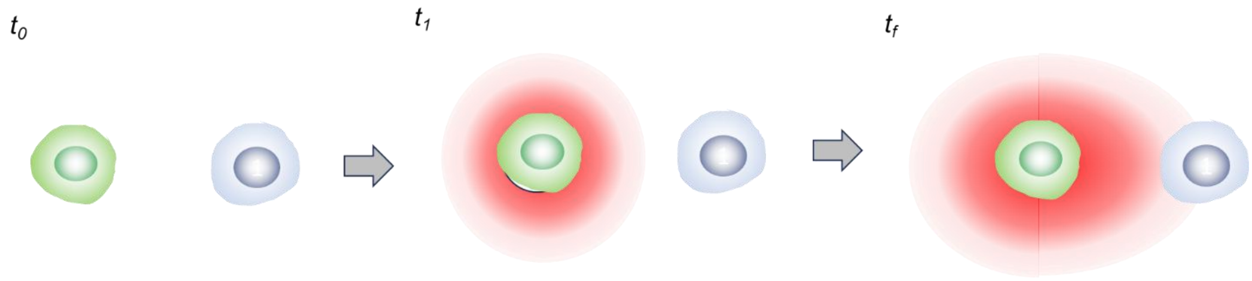


**Figure S11.** Schematic showing polarization of secreted cytokine during cell-cell communication
